# Integrative Spatial Modelling of Cellular Plasticity using Graph Neural Networks and Geostatistics

**DOI:** 10.1101/2025.09.24.678189

**Authors:** Eloise Withnell, Cenk Celik, Maria Secrier

**Affiliations:** UCL Genetics Institute, Department of Genetics, Evolution and Environment, University College London, Gower Street, London, WC1E 6BT, UK

## Abstract

Cellular plasticity - the ability of cells to change phenotype in response to intrinsic and environmental cues - is central to development, regeneration, and disease, but remains difficult to quantify due to its dynamic, context-dependent nature. Here we introduce a framework that unites AI and geostatistics - graph neural networks and spatial regression models - to both predict and explain cell state variation in spatial transcriptomics data. We formalize state predictability as a quantitative proxy for plasticity, where stable states are predictable and plastic states are not. Applied to epithelial-mesenchymal plasticity (EMP) in breast cancer, the framework shows that mesenchymal states are stabilized by recurrent copy number alterations and microenvironmental cues, whereas hybrid states remain unpredictable and plastic. It also uncovers a long-range influence of myofibroblasts on EMP, demonstrating that stromal remodeling can propagate plasticity across tissue regions. Our framework yields interpretable, scale-aware insights into intrinsic and extrinsic drivers of state transitions, providing a broadly applicable strategy for modelling dynamic cell states in spatial biology.

## Introduction

Cell plasticity, the ability of cells to alter their phenotype in response to intrinsic and environmental stress^1^, is a defining property of multicellular life. It underpins embryonic development, tissue regeneration and immune responses, but also fuels pathological processes such as cancer progression and therapy resistance^2^. A hallmark example is epithelial-mesenchymal plasticity (EMP), whereby epithelial cancer cells adopt mesenchymal traits, facilitating therapy evasion and survival in adverse tumor microenvironments (TMEs)^3,4^. Despite its importance, plasticity remains difficult to quantify and predict because it is highly context-specific: mutations and epigenetic remodelling may increase a cell’s responsiveness to environmental cues^5,6^, while the cellular microenvironment exerts spatially structured pressures through gradients of nutrients, oxygen, matrix stiffness and paracrine signalling^7–9^, resulting in heterogeneous and often reversible phenotypic shifts^10,11^. Understanding what drives these shifts in cells could allow us to manipulate them towards more targetable disease states.

Experimental systems such as cell lines or organoids have provided insights into drivers of plasticity, e.g. through lineage tracing or FACS sorting^8,12–14^, but often lack the spatial resolution or scalability to assess how multiple intrinsic and extrinsic factors jointly shape cell states. Computational approaches such as PATH, which infers phenotype heritability from genetic distances^15^, or chromatin-state prediction models using prediction uncertainty as a proxy for plasticity^16^, demonstrate the potential of quantitative approaches but are not designed to account for spatial structure. Methods like CellCharter^17^ or SIMVI^18^ leverage spatial transcriptomics to infer cell intrinsic and spatially-induced variation within the tissue but lack interpretable coefficients, making it difficult to quantify the influence of individual cell types or intrinsic factors such as copy number alterations. Tools like GASTON^19^ or CRAWDAD^20^ quantify spatial gradients and distance-dependent cellular relationships, underscoring the importance of modelling spatial scale explicitly^21^. Our method, SpottedPy^9^, applies geospatial statistics to map plastic niches and their neighborhood structure at variable scales, but like other methods^22,23^, cannot jointly quantify intrinsic and extrinsic effects in an interpretable way - highlighting the need for integrative models with both predictive capacity and transparency.

To address this, we draw inspiration from AI-based and ecological modelling frameworks, both of which offer tools to formalize genotype-environment–phenotype relationships. Graph neural networks (GNNs) capture structured dependencies and have been applied in spatial transcriptomics to model intercellular communication, tissue architecture, and predict cell-state transitions^24–26^. The challenge, however, is not only to predict but also to interpret. In ecology and geostatistics, spatial models quantify the influence and scale of different variables on species traits. For instance, spatial autoregressive models of tropical forests quantify how much variation in leaf and wood traits comes from genetics versus external factors like soil fertility, water availability, and neighbour effects^27^. Analogous models could be envisioned to disentangle intrinsic from extrinsic pressures within cellular niches, as previously proposed in related contexts^28–31^.

Here, we propose an interpretable, multi-layered modelling approach designed to quantify the impact of intrinsic (genomic) and extrinsic (microenvironmental) factors on cellular plasticity from spatial transcriptomics. We combine Graph Neural Networks (GNNs)^32^, Spatial Error Models (SEMs)^33^, and Geographically Weighted Regression (GWR)^34,35^, which offer complementary strengths: GNNs uncover nonlinear, higher-order cellular interactions, while geostatistical models provide a statistical measurement of spatial heterogeneity, enabling quantification of short- and long-range influences on specific cells. This integration enables not only state-of-the-art predictive accuracy, but also interpretable, scale-aware coefficients that disentangle genomic from extrinsic microenvironmental drivers of cell behaviour.

As a proof of concept, we apply the framework to epithelial–mesenchymal plasticity (EMP) in breast cancer, showing that mesenchymal states are the most predictable (stable), while hybrid states are the least predictable (plastic), with myofibroblasts exerting a long-range influence on these transitions. In parallel, benchmarking on mouse cortex data illustrates how the framework generalizes to developmental systems. Together, these applications demonstrate how integrative AI–geostatistical modelling can yield transparent, quantitative insights into the spatial determinants of cellular plasticity.

## Results

### A framework to quantify cell plasticity from spatial data

We propose a general framework to model how intrinsic and extrinsic pressures influence cell states and plasticity potential (**Fig.1**). We hypothesize that cell states which are difficult to predict from genomic or experimental context exhibit greater plasticity potential, reflecting dynamic and adaptable behaviour. In contrast, predictable states likely represent more stable phenotypes, constrained by their microenvironment. To test this, we applied three complementary models— GNNs, SEMs, and GWR—to model EMP in breast cancer spatial transcriptomics data (**Fig.1**). We approximate intrinsic effects on cancer cell state through copy number alterations (CNAs) and extrinsic effects through interactions with other cells in the TME. If the epithelial-to-mesenchymal transition (EMT) cannot be reliably predicted based on features of the TME, then the spatial positioning of cancer cells within the TME does not impact this process. If we cannot predict EMT using genomic factors, then these do not impact the transition.

**Figure 1.**
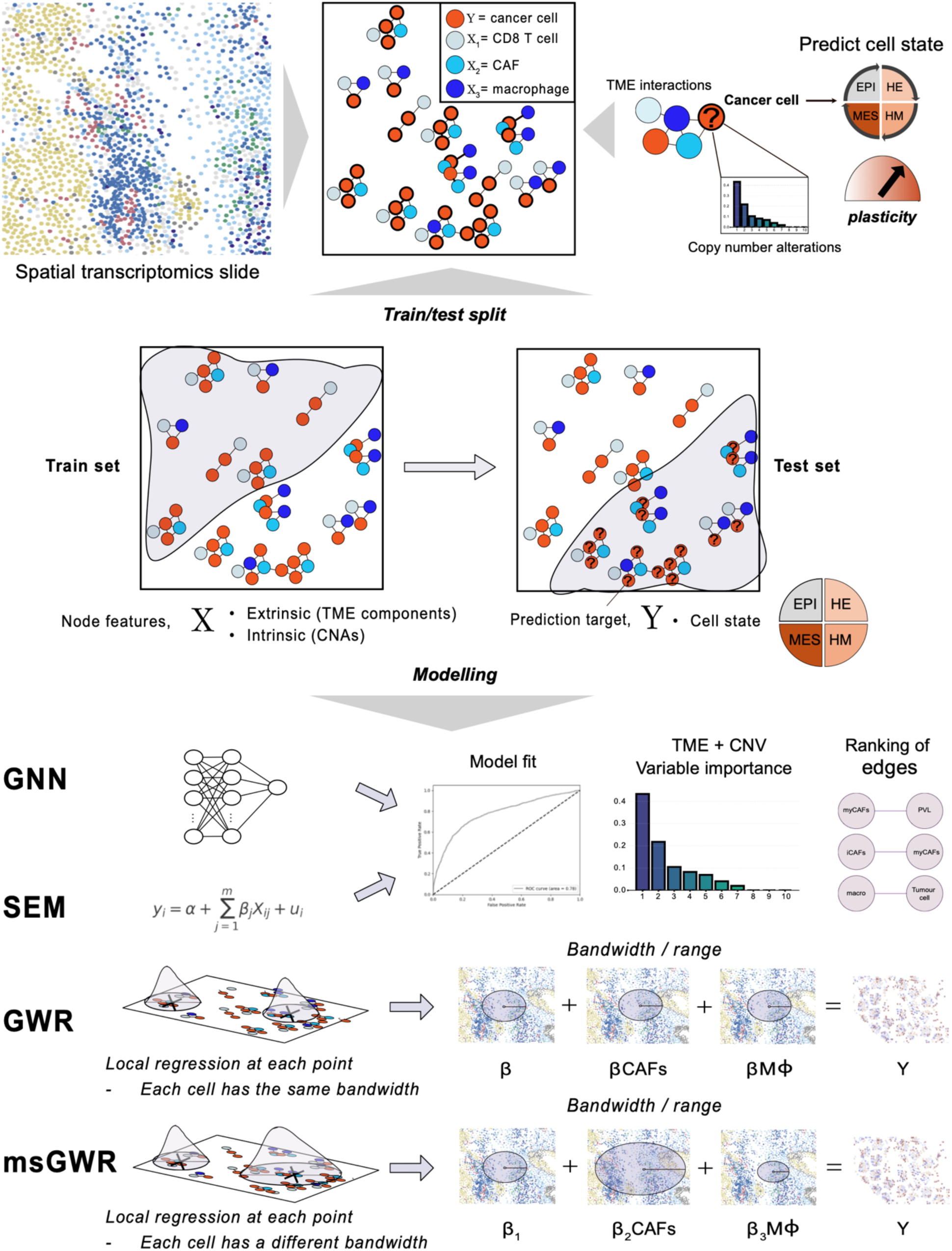
Overview of the computational framework to model cell plasticity in spatial transcriptomic data. The framework is exemplified using EMP, an example of cancer cell plasticity. **Top panel:** Cancer cells and their interactions with other cell populations in the TME are mapped from spatial transcriptomics data. EMP states of cancer cells (EPI: epithelial; HE: hybrid-epithelial; HM: hybrid-mesenchymal; M: mesenchymal) are modelled based on extrinsic (tumour microenvironment, TME) and intrinsic (copy number alterations, CNAs) variables. **Middle panel:** Training is performed on part of the spatial transcriptomics slide, with testing performed on the remaining part. **Bottom panel:** Different modelling approaches (GNN – graph neural networks; SEM – spatial error modelling; GWR – geographically weighted regression; msGWR – multiscale geographically weighted regression) are used to capture intrinsic and extrinsic effects on cell plasticity (Y). Both GNNs and SEMs offer measures of model fit, variable importance ranking and ranking of edges corresponding to cell-cell interactions. GWR and msGWR allow the modelling of a cell’s impact on another cell with fixed (GWR) or variable (msGWR) bandwidths (or ranges) of influence. CAFs = cancer-associated fibroblasts, MΦ = macrophages.

**Figure 2.**
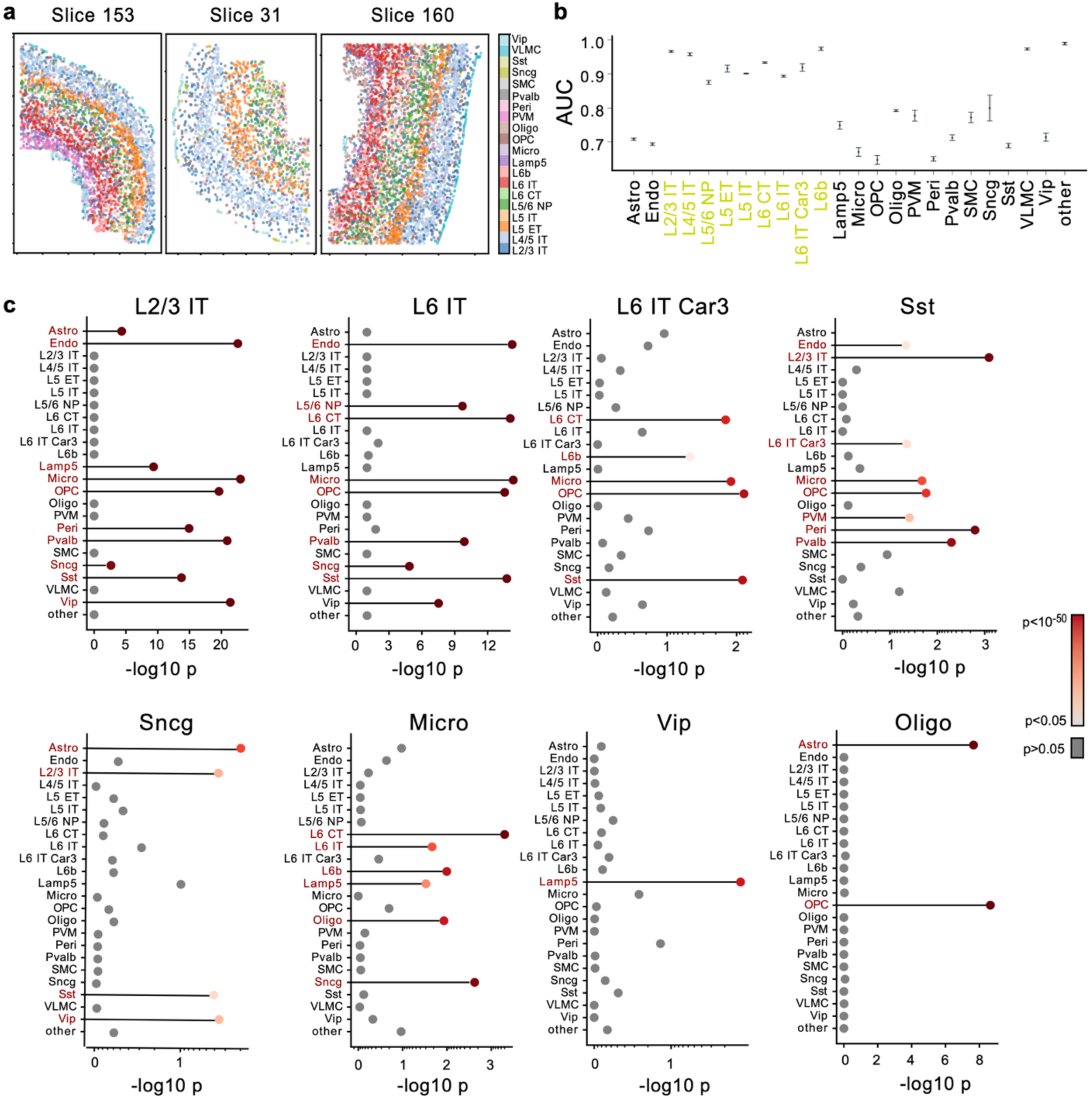
Graph neural networks capture spatially-defined cell identities in the mouse cortex. **a.** MERFISH cell types visualised on distinct mouse cortex slices. **b.** GNN prediction performance for each cell type in the MERFISH dataset (quantified as AUC scores). Spatially constrained cells are highlighted in yellow. **c.** Node importance values for selected cell types. Significantly positioned nodes with respect to the predicted cell type are highlighted in red.

**Figure 3.**
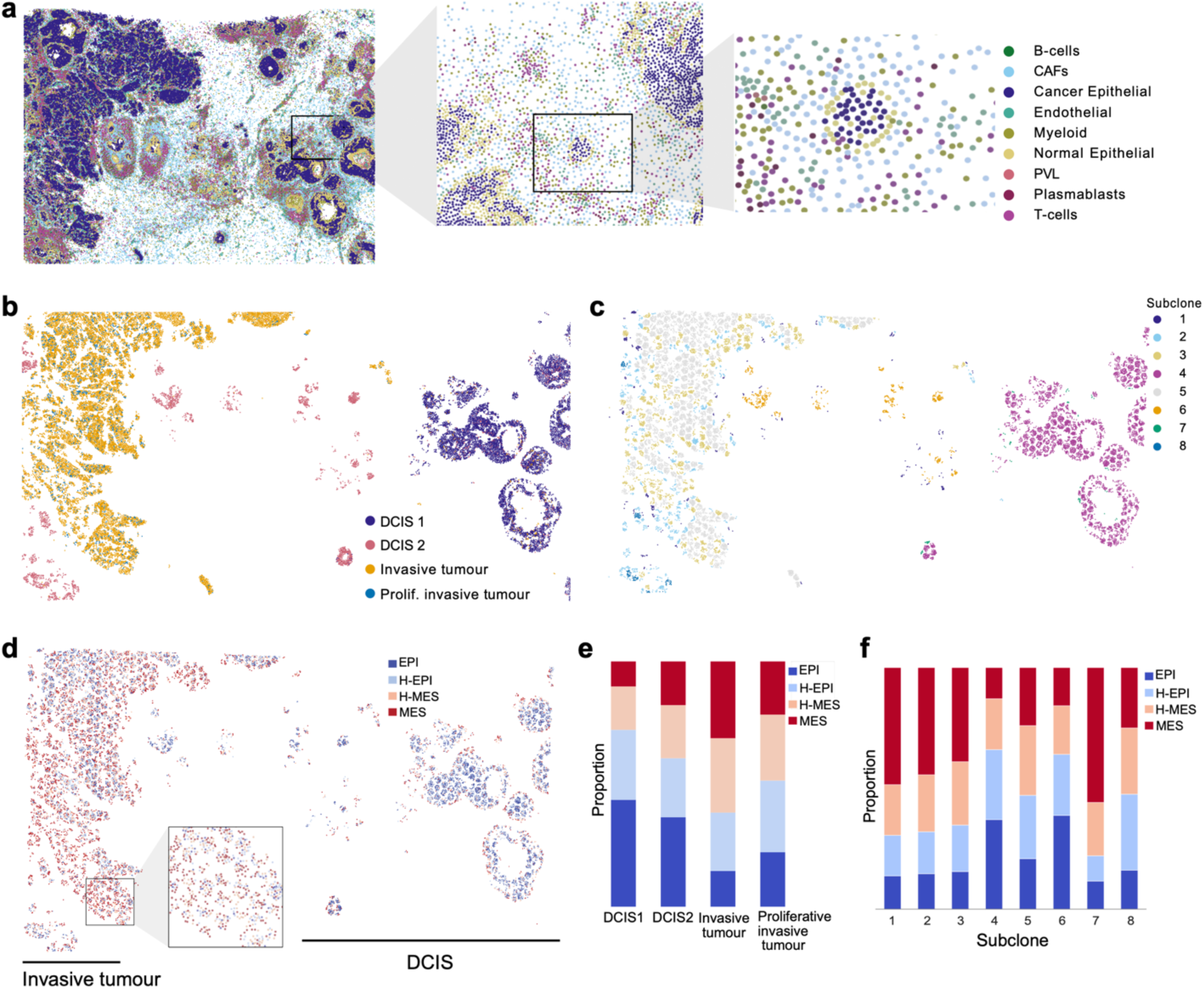
Clonal evolution and EMP state mapping in breast cancer Xenium data. **a.** Cell type annotations labelled in the Xenium breast cancer dataset. **b.** Pathologist annotations visualised across the Xenium dataset. DCIS = ductal carcinoma in situ. **c.** Tumour subclones identified by SCEVAN, visualised across the Xenium slide. **d.** EMP states in the cancer cells identified in the Xenium dataset, mapped across the slide. EPI = epithelial, H-EPI = hybrid-epithelial; H-MES = hybrid-mesenchymal; MES = mesenchymal. **e.** EMP state distribution across the regions annotated by the pathologist. **f.** EMP state distribution across the tumour subclones.

**Figure 4.**
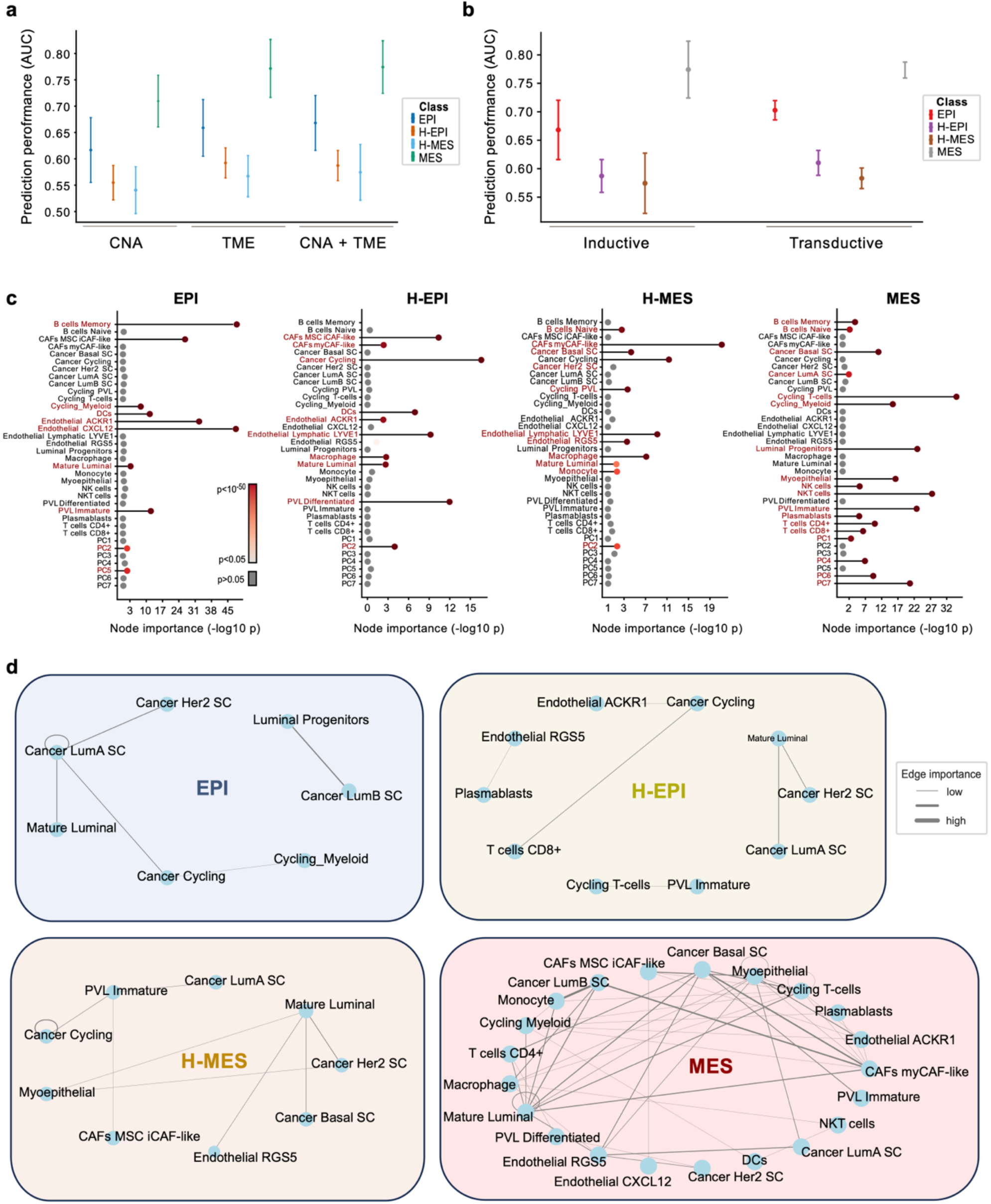
Graph neural networks model intrinsic and extrinsic effects on cellular plasticity in breast cancer. **a.** Predictive accuracy for EMP states based only on CNAs, only on the TME and combining both factors of influence. AUC ranges from 10-fold cross-validation are plotted with minimum, maximum and mean values marked. Different colours denote different predicted plastic states. **b.** Predictive accuracy using different methods of training (inductive/transductive). Annotations as in (a). **c.** Node importance values for each EMP state. Significantly positioned nodes are highlighted in red. **d.** Network graphs for each EMP state.

**Figure 5.**
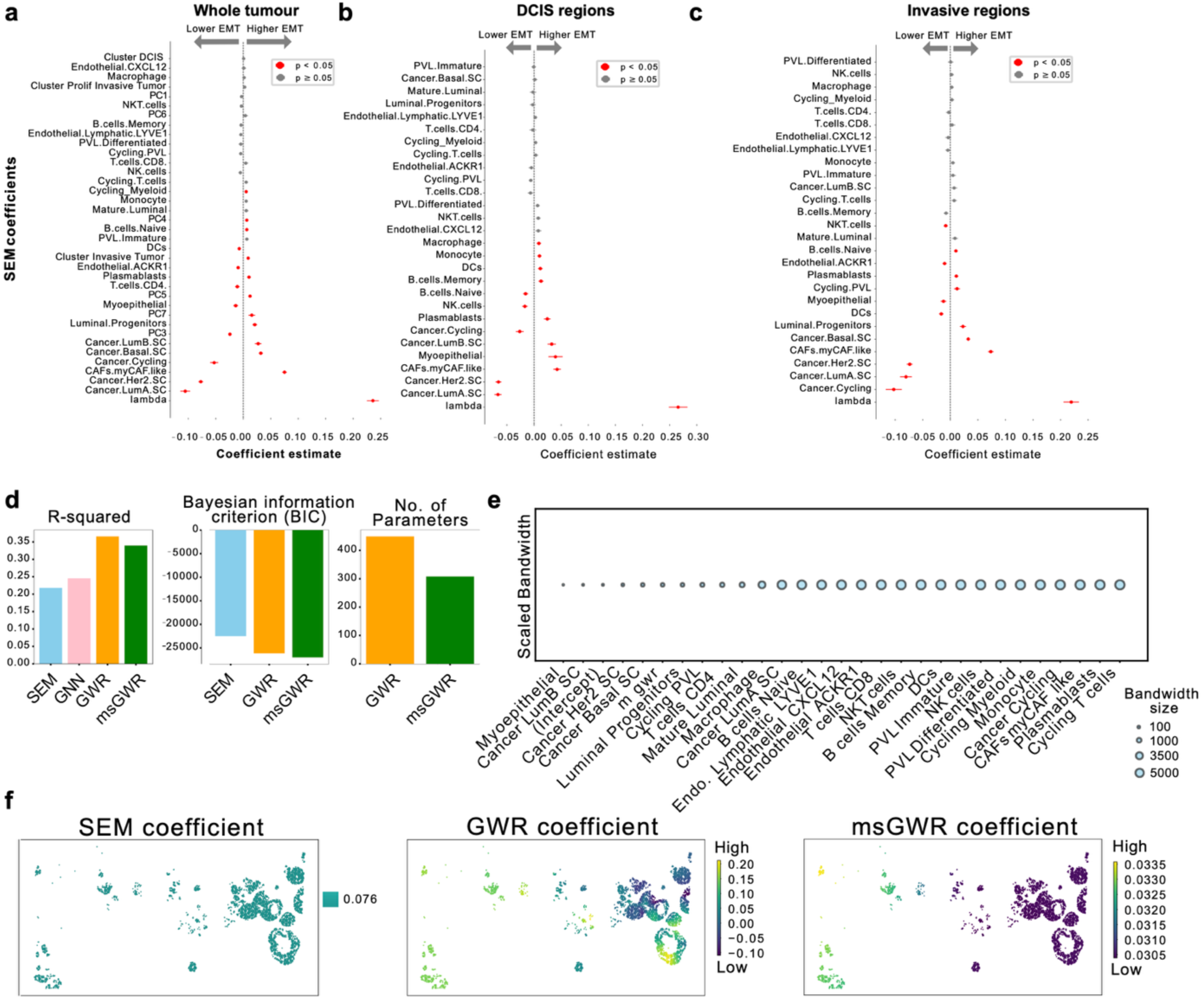
Spatial regression quantifies the impact and range of action of various intrinsic and extrinsic influences on cancer cell plasticity. **a.** Estimated variable importance outputted by the SEM regression model EMT based on data from the breast cancer Xenium slide. **b.** The same as (a) but for DCIS regions only. **c.** The same as (a) but for invasive regions only. **d.** Metric comparison for SEM, GNN, GWR and msGWR models, including R^2^, BIC and number of parameters employed in the models. **e.** Bandwidths for each variable used in the msGWR model. Lower bandwidth indicates a shorter (local) range of influence of various cells on cancer cells undergoing EMT. **f.** Visualisation of the SEM, GWR and msGWR coefficients for CD8+ T-cells across the breast cancer slide. Higher coefficients indicate higher CD8+ T cell influence. Going from the right (DCIS) towards the left (more invasive) part of the image, we see the expected increased CD8+ T cell association with EMT most accurately captured with the msGWR model.

First, we apply GNNs to predict whether the cancer cell is in an epithelial, hybrid or mesenchymal state based on its genomic changes and the TME. By interpreting the importance of nodes and edges in the GNN predictions, we gain insights into the model’s learning process and understand the cellular networks important for this transition. We then use spatial regression to further characterize EMP as a continuous process and show how msGWR can quantify the range of influence different immune or stromal cells have on the cancer cells, as shown in the next sections.

### Modelling spatially constrained states using GNNs

GNNs are uniquely suited to modelling spatial interactions, as they learn from relationships between neighboring nodes—here, cell types—without relying solely on distance-based features. To illustrate the concept of capturing spatial effects on cell identity, we first benchmarked the method on a MERFISH mouse cortex dataset containing spatially ordered and unordered cell types^36^ (**Fig.2a**). Layered neuronal subtypes, such as layer 2/3 and 4/5 intratelencephalic neurons (L2/3 IT, L4/5 IT), layer 5 extratelencephalic neurons (L5 ET), layer 5/6 near-projecting neurons (L5/6 NP), layer 6 corticothalamic neurons (L6 CT), layer 6b neurons (L6b), and vascular leptomeningeal cells (VLMC) exhibit well-defined distinct spatial organisation within the motor cortex, each with distinct roles in cortical processing^37,38^ (**Fig.2a**). In contrast, more plastic cells like pericytes, which can differentiate into smooth muscle cells, fibroblasts or osteoblasts in response to injury, inflammation or changes in blood flow^39–41^, or oligodendrocyte precursor cells (OPCs), which differentiate into oligodendrocytes in response to local myelination demands^42^, display more variable distributions.

We hypothesized that spatially constrained cell types would be more accurately predicted by the GNN model than more plastic or randomly distributed ones. To test this, we trained the model on a subset of the MERFISH slides and evaluated performance on independent slides to avoid data leakage, averaging results across 10 distinct train-test splits (Methods, **Supplementary Table 1**). For each experiment, we masked a specific cell type (used as the prediction label) and provided only the remaining cell types as input. Consistent with our hypothesis, spatially constrained cell types such as L2/3 IT, L4/5 IT, L5 ET, L5 IT, L5/6NP, L6CT and VLMCs were predicted with high accuracy, achieving AUC values above 0.85 (**Fig.2b**). In contrast, OPCs and pericytes, known to present higher plasticity inherently linked to more variable locations and functions, were amongst the least predictive. This supports the idea that **poorly predictable states reflect higher plasticity**.

Beyond prediction, the GNN model allowed us to interpret the learned spatial relationships (**Fig.2c**, **Supplementary Fig.1**). Node and edge importance analysis revealed biologically expected associations. For instance, L2/L3 IT neurons were influenced by VIP interneurons, microglia and endothelial cells, consistent with known regulatory interactions and vascular support mechanisms^43–45^ within the same cortical layer^46^. L6 IT and Car3 neurons were positionally linked to L6 CT neurons and Sst interneurons, both known to form local circuits and synaptic connections^47–49^. VIP and Lamp5 neurons, enriched in superficial layers (1-3), also exhibited an expected spatial association^50^. Finally, the association between oligodendrocytes and OPCs reflected their expected differentiation trajectory^51^.

Thus, GNNs can both capture and explain spatial dependencies of cells with different identities and functions, establishing a foundation for modelling plasticity through predictability.

### Modelling cancer cell plasticity using GNNs

Having confirmed the model aligns with biological expectations, we next applied GNNs to interrogate EMP in a Xenium breast cancer spatial transcriptomics dataset from Janesick et al^52^, consisting of nine major cell types including cancer epithelial cells, cancer-associated fibroblasts (CAFs), immune, endothelial and stromal populations (**Fig.3a**). We also integrated pathologist annotations of ductal carcinoma in situ (DCIS) and invasive regions within the tissue (**Fig.3b)**. To model intrinsic factors, we inferred copy number alterations (CNAs) in cancer cells using SCEVAN^53^ and identified eight tumour subclones displaying alterations frequent in breast cancer such as 17q22-24 amplification^54^, 8q24.3 amplification spanning the *MYC* oncogene^55^, or 11q13.4-q25 deletion containing cyclin D1^56^ (**Fig.3c**, **Supplementary Fig.2a-b**). To facilitate downstream modelling, we performed principal component analysis (PCA) on the CNA matrix (**Supplementary Fig.2c**), extracting PC loadings that mirrored phylogenetic relationships among subclones (**Supplementary Fig.2d**) and captured key events in breast carcinogenesis such as *CDKN2A and BRCA2* deletions (PC1), *ATM* deletions (PC4), *EGR1* gains or *ERBB2* amplifications (PC7) (**Supplementary Table 2**).

We then assessed the EMT state of the cancer epithelial cells using a 195-gene hallmark EMT signature from MsigDB (see Methods). In light of recently reported challenges in capturing signatures with targeted ST gene panels^57^, we ensured that the reduced gene coverage in Xenium (20 EMT genes) retained strong discriminatory power compared to a matched Visium slide (Methods, **Supplementary Fig.3a**). Although the number of EMT states in breast cancer remains debated, more than one hybrid state has been suggested^58–60^. In line with this, we took a parsimonious approach where we defined four EMT states (epithelial, epithelial-hybrid, hybrid-mesenchymal, mesenchymal) by binning the EMT scores into quartiles (**Fig.3d**). Regardless of the precise number of states, this classification serves as a proof of concept for modelling cell plasticity in spatial data. Because CAFs and mesenchymal tumour cells share transcriptional features^61,62^, we removed potentially misclassified mesenchymal cancer cells with high expression of *LUM*, a discriminative marker for CAFs which we inferred from single cell RNA-seq data (Methods, **Supplementary Fig.3b-h**). The resulting EMT state annotations (**Fig.3d**) reflected the expected transformation from the DCIS regions dominated by epithelial, non-invasive tumour cells to the invasive margin enriched in mesenchymal cells (**Fig.3e**), which were also subclonally segregated (**Fig.3f**).

We next trained a GNN to predict EMT states in cancer cells based on intrinsic information (CNAs) alone, the TME (extrinsic) information alone or combined. The mesenchymal state was the most predictable, with 71% mean AUC based on CNAs alone, rising to 77% when TME influence was considered (**Fig.4a, Supplementary Table 3a-b**). Combining CNA and TME factors did not further improve the performance of the model (**Fig.4a, Supplementary Table 3c**), indicating that while the microenvironment is the primary determinant of EMT (as expected^63^), specific genomic alterations may be selected to stabilize plastic shifts. Unsurprisingly, hybrid states were most challenging to predict (**Fig.4a, Supplementary Table 3a-c**), in line with their increased plasticity. Thus, GNNs can serve both to identify drivers of a specific cancer cell phenotype and to assess plasticity more generally, where low AUCs suggest unstable states with high plasticity potential. These observations were consistent regardless of whether the dataset was split for training and testing through inductive graph learning, based on separate spatial splits, or transductive learning, via random node masking (**Fig.4b**, **Supplementary Fig.4a, Supplementary Table 3d-f**).

The mesenchymal state was most strongly associated with cycling T cells and NKT cells, suggesting an immune active state (**Fig.4c**). It was also impacted by CNAs pertaining to PC6 (including *CD44* amplification on chromosome 11, a classical EMT event^64^) and PC7 (including *ERBBR2* amplification on chromosome 17, also linked with EMT^65^) (**Supplementary Table 2**). Hybrid mesenchymal cells presented immune evasion features, with myCAFs and macrophages forming the most important nodes. Epithelial cancer cells were most strongly associated with memory B cells, *CXCL12*+ and *ACKR1*+ endothelial cells, indicative of chemokine signalling and adaptive immune responses. These endothelial cells differed from the *LYVE*+ and *RGS5*+ cells linked to the hybrid states, which have been implicated in metastasis, invasion and hypoxia response^66,67^. Similar associations were found when using a simplified two-state EMT model (**Supplementary Fig.4b**), reinforcing the stability of overall cell type trends and the accuracy of the GNN explanations. This simplified model also demonstrates the advantage of a four-state model in providing additional granularity into the EMT process.

We observed progressive rewiring and co-opting of TME factors as the cancer cells progressed from epithelial through hybrid and towards mesenchymal states (**Fig.4d**). While the epithelial state was mostly defined by luminal and cycling cancer cell interactions, consistent with an epithelial cancer cell core that is less exposed to the TME^68^, a progression towards basal phenotypes and increased co-option of endothelial cells and iCAFs was observed in the hybrid states, while the mesenchymal state displayed the highest dependence on the TME.

We also tested the GNN approach on a continuous EMT score (**Supplementary Fig.4c-d**). The TME explained double the variance in EMT score compared to CNAs, aligning with the idea that EMT can be modulated rapidly in response to environmental conditions^63^. In contrast, stable, inherited genomic changes appear to associate with large jumps in the EMT landscape towards more stable transitions i.e. shifting between fully epithelial and fully mesenchymal states. These transitions may require more profound cellular reprogramming, which aligns with the longer timescales of clonal evolution compared to the immediate, dynamic nature of TME interactions.

### Modelling cancer cell plasticity using spatial regression

While GNNs provide valuable insights into spatial dependencies by modelling the graph structure, they are limited in accounting for confounding effects or spatial heterogeneity. Geostatistical models address this by identifying context-specific spatial effects, such as those arising in different regions of the tissue or due to different cancer subtypes^69^. Here, we apply spatial error models (SEMs) - a type of spatial regression that incorporates a spatial error term to account for unobserved spatial heterogeneity^89^. This allows us to estimate conditional dependencies between cells while accounting for spatial autocorrelation^31^.

When applying this method, we observed that the TME explained a larger fraction of the variance than CNAs (**Supplementary Fig.5a**), consistent with GNN results. myCAFs emerged as the TME cell type with the greatest spatial influence on EMT (**Fig.5a**), as previously reported by us^9^. CNAs remained significant when the effects of the TME were regressed out, suggesting that certain genomic alterations that become fixed during cancer evolution help push cancer cells into an EMT state, independent of colocalization with other cells in the TME. The spatial error coefficient (λ) was significant in all SEMs, underscoring the relevance of spatial effects. Additionally, by regressing out the effects of DCIS versus invasive tumour, we found that invasive regions were significantly associated with a mesenchymal cancer state, as expected (**Fig.5a**). Separate SEMs for ductal and invasive regions (**Fig.5b-c**) revealed consistent myCAF associations with a mesenchymal state, but also regional differences: for instance, myoepithelial cell presence was linked with higher EMT in DCIS areas, but lower EMT in invasive regions, in line with their dual role in maintaining epithelial integrity but also promoting tumour progression through TGFβ signalling^70,71^.

It is important to note that whilst SEMs could provide additional insights that GNNs cannot, when comparing the overall regression metrics GNNs outperform SEMs (**Supplementary Fig.5b**).

The varying influence of myoepithelial cells across regions highlights spatial heterogeneity (also known as spatial non-stationarity). Most methods assume uniform spatial relationships, which rarely reflect biological reality. Geographically weighted regression (GWR)^34^ allows us to assess spatial heterogeneity by asking whether immune or stromal cells exert different influence on cancer cells depending on their proximity. The key advantage lies in moving beyond global averages to assess *local* short-range and long-range relationships. While classical GWR assumes all cells influence each other over the same spatial scale, multiscale GWR (msGWR) assigns each cell its own bandwidth, reflecting its range of influence^35^ (**Fig.1**). Larger bandwidths indicate broader influence, smaller ones more localised effects. Knowing this could inform whether therapeutic interventions should be local or systemic.

When applied to the data, GWR-based models showed improved performance over GNNs and SEMs (**Fig.5d**). Although msGWR did not outperform GWR, it did reduce model complexity, as reflected by lower BIC value and fewer parameters (**Supplementary Table 4**). Thus, while short-range interactions can be sufficient for assessing EMP, msGWR offers the added advantage of revealing the spatial scale at which each cell operates (**Supplementary Table 5**). This analysis confirmed that myoepithelial cells exert short-range influence (**Fig.5e**), consistent with the SEM modelling. In contrast, myCAFs displayed a longer-range influence on EMT, likely due to their role in ECM deposition, remodelling and tissue stiffening, which may promote widespread EMT^72^.

Finally, the msGWR approach enables precise spatial mapping of the range of influence of selected cell populations. For instance, compared to SEM or GWR, the flexible range of action allows us to capture local associations between CD8+ T cells and cancer cells undergoing EMT that are more in line with an expected progressive CD8+ T cell enrichment along the DCIS-to-invasive cancer cell transformation gradient (**Fig.5f**). This underscores the value of local over global modelling in ST data.

## Discussion

We present an analytical framework that allows the precise spatial modelling of intrinsic and extrinsic factors contributing to cellular plasticity, which can be flexibly tuned to interrogate various phenotypes and assess the stability of cell states. Using it to explore EMP in breast cancer, we show that predictability of cell state from context could serve as a proxy for plasticity: mesenchymal states were most predictable based on environmental and genomic factors, consistent with a more deterministic, stable phenotype within a remodelled niche that becomes locked in through clonal evolution, whereas hybrid states were least predictable, indicating higher plasticity. These findings point to a model where hybrid states function as highly adaptable intermediates that can rapidly rewire their dependencies on the surrounding microenvironment, while mesenchymal states consolidate into more stable attractors of the EMT landscape. Moreover, the discovery that myCAFs exert long-range influence on EMT suggests that stromal remodeling can propagate plasticity across tissue regions, which could open novel therapeutic angles that extend beyond immediate cell–cell contacts.

This study serves as a **proof of concept** for how AI and geospatial modelling could be employed to quantify cellular plasticity. GNNs revealed spatial relationships and non-linear dependencies between cell states and their environment, while geostatistical models—particularly msGWR— offered interpretable, scale-aware insights into local and long-range influences. Although GNNs outperformed spatial regression models in predictive accuracy, SEM and GWR provided more interpretable coefficients and allowed for statistical control of molecular confounders. Combining these tools enables both **prediction and explanation** of spatial determinants of plasticity across scales.

We propose prediction scores (e.g. AUC values from the GNN model) could serve as a quantitative metric for plasticity potential, reflecting the extent to which cellular phenotypes adapt to TME pressures or intrinsic genomic instability. Further enhancements of such models could involve entropy-based metrics, such as those used by Burdziak et al^16^, where higher entropy across classes would suggest increased plasticity.

To enhance interpretability, we simplified EMP into four discrete states using a binning approach. More nuanced strategies like trajectory inference^73–76^ or Gaussian mixture modelling^77,78^ could be used in the future to refine EMT state classification. To model intrinsic factors, we used CNAs inferred from spatial transcriptomics and PCA to manage multicollinearity, which limits interpretability at the gene level. Future versions should incorporate single-cell genomic and epigenetic data, and explore alternative strategies for modelling genomic variation. Moreover,

TME features were represented at the cell level, but capturing the full spectrum of EMT drivers likely requires modelling additional layers within the tissue, such as chemokine gradients, matrix stiffness and hypoxia. We see this framework as a foundational step toward building more comprehensive models of cell plasticity across systems.

To further improve predictions, future methods may integrate heterogeneity-aware deep learning spatial models^79^ or graph transformers^80^, which can mitigate GNN over-smoothing^81^. Causal modelling^82^ could clarify directional effects between TME, mutations and plasticity. Recent methods like SIMVI^18^ offer promising metrics to disentangle intrinsic and spatial effects; while not yet fully interpretable, they could offer additional insights into the balance between intrinsic and extrinsic factors driving cell plasticity. While this study focused on conceptually probing the combined power of AI and geostatistics in assessing cellular plasticity, future work should validate findings in larger datasets that will allow to inspect intra-patient heterogeneity. Furthermore, determining causal effects would require additional modelling or experimental validation through perturbation studies.

The framework proposed here enables spatially-resolved, quantitative insights into the drivers of cellular plasticity, and is broadly applicable across developmental, regenerative, and pathological contexts. Ultimately, such integrative spatial modelling approaches could offer a foundation for identifying phenotypic states—and transitions between them—that could be targeted or stabilized therapeutically.

## Methods

### MERFISH dataset processing

A mouse motor cortex MERFISH dataset was downloaded from Zhang et al^36^. The data consists of 61 tissue slices and 280,000 cells. The cells were annotated using 258 genes in the original study. A total of 23 cell subclasses were identified. We used the coordinates and cell type annotations as provided by the authors.

### Xenium breast cancer dataset processing

A Xenium breast cancer dataset consisting of 167,780 cells and 307 genes and a matched Visium dataset consisting of 4992 spots and 18,085 genes was obtained from Janesick et al^52^. The sample was Stage II-B, ER + /PR − /HER2 + formalin-fixed paraffin-embedded (FFPE) breast cancer tissue. To align the Visium data to the Xenium data, the SpatialData^83^ Python package was used. SpatialData uses landmark points in the images to transform data into a common coordinate system. Cell annotations were used as calculated in Marconato et al^83^. Additional subtype annotations as labelled by a pathologist (including whether the region is invasive versus DCIS) were used as described in Janesick et al^52^.

We used Scanpy^84^ for pre-processing, using default parameters. Specifically, we filtered out genes that were in less than 5 cells, and ensured each cell had a minimum of 75 counts. For Visium we ensured the mitochondrial fraction was less than 15%, the number of genes was larger than 500 and cells had a minimum gene fraction of 0.2.

### Tumour clonal reconstruction

To perform subclonality estimation in cancer cells, we utilised the matched Visium dataset due to its ability to estimate copy number amplification, currently not possible with Xenium data because of its limited gene coverage, which limits the detection of copy number alterations. Using SCEVAN^53^, a fast variational algorithm using multichannel segmentation, we added further evidence for the tumour cells, which had been previously identified using Cell2location^85^. We chose SCEVAN as it does not require user provided parameters like other similar methods require, and instead automatically estimates the highly confident normal cells based on count data to use as a baseline. Using SCEVAN, we identified subclones and estimated the chromosomal regions affected by alterations within each subclone.

### PCA decomposition of CNAs

To reduce the dimensions of the sub-clonal alteration matrix returned by SCEVAN (8 subclones and 160 altered regions) we ran principal component analysis (PCA) on the dataset. We obtained principal components capturing the main sources of variation within the dataset. This allowed us to derive principal components that encapsulate the key sources of copy number variation within the data.

### EMT annotation

We used scanpy.score_genes to score the EMT hallmark gene signature from MSigDB^86^ on the Xenium dataset. We ascertained a high correlation between the set of EMT genes found in Xenium and the Visium datasets, to ensure that the limited gene coverage found in Xenium did not impact the gene signature scoring. To obtain states of epithelial, hybrid and mesenchymal cancer cells, we binned the EMT hallmarks score into four quartiles, representing an epithelial state, an epithelial-hybrid state, a mesenchymal-hybrid state and a mesenchymal state.

### Marker genes specific to EMT tumour cells

We performed further quality control to minimize mislabeling between myCAFs (which display mesenchymal features and an enrichment of EMT genes^61^) and mesenchymal cancer cells. To this end, we analysed key differentiating genes in a well annotated breast cancer atlas scRNA-seq dataset from Wu et al^87^. In this dataset, we filtered for only the subset of genes that were present in Xenium. We have previously annotated this dataset with EMT states. We used sc.tl.rank_genes_groups within the scanpy package to find the top marker genes differentiating between the labelled mesenchymal cancer epithelial cells and CAFs (**Supplementary Fig. 3b**). We selected the top 10 markers (**Supplementary Fig. 3c**) and employed Cohen’s d score to quantify the effect size and discriminative power of each candidate marker in distinguishing cancer cells from CAFs (**Supplementary Fig. 3d**). Cohen’s d measures the standardised difference between the means of two groups, in this case, the expression levels in CAFs versus mesenchymal cancer cells, and identified the number of genes required to accurately distinguish between the cell types. *LUM* emerged as the most discriminative gene (Cohen’s *d* = –4.26), outperforming all other combinations (Cohen’s *d* ranging from –3.45 to –3.62) (**Supplementary Fig. 3d-f**). The *LUM* gene encodes lumican, a small leucine-rich proteoglycan involved in collagen fibrillogenesis and extracellular matrix organisation, and has been implicated in regulating cell migration, proliferation and tumour progression^88^. We then investigated *LUM* expression in the Xenium data and filtered out mesenchymal-labelled cancer cells with abnormally high *LUM* expression i.e. higher than the 1st quartile range of *LUM* expression for CAFs (**Supplementary Figure 3g-h**). 138,780 cancer cells remained after filtering.

### GNN approach

To construct the GNN prediction approach, we adapted the widely used Graph Convolutional Network (GCN) framework.

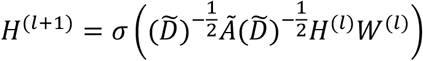

where, *H*^*(l)*^ is the matrix of node feature representations at layer *l*, with *H*^*(o)*^ = *X*; *H^(l+1)^* is the matrix of node feature representations at layer *l* + 1; *Ã* is the adjacency matrix with added self-loops, i.e., *Ã* = *A* + 1; *D̃* is the degree matrix of *Ã*; *W^(l)^* is the learnable weight matrix for layer *l*; and *σ* is the activation function (e.g. ReLU).

The datasets were loaded into a PyTorch Geometric dataset, where we adapted custom classes to include various features and configurations tailored to our experiments. These configurations included incorporating cell type and/or copy number information, as well as modifying the training paradigm to be either inductive or transductive. On the MERFISH dataset, we utilised a graph neural network (GNN) with three graph convolutional layers, a learning rate of 0.01, and trained the model for 200 epochs. A dropout rate of 0.5 was applied to regularise the model and prevent overfitting. Similarly, for the Xenium breast cancer dataset, we trained a GNN with the same architectural configuration, three layers, a 0.01 learning rate, and 200 epochs, with a dropout rate of 0.5. We also used Graph Attention Networks (GAT) but found no difference in accuracy. The SquidPy (Spatial Single-Cell Analysis in Python) package^89^ was used for graph construction using sq.gr.spatial_neighbors and NetworkX was used for further graph manipulation.

Mean squared error (MSE) loss was used for continuous training, while cross-entropy loss was used for classification tasks involving categorical labels. This enabled the implementation of both GNN regression and GNN classification models. We adjusted for spatial autocorrelation by calculating the Moran’s I statistic and propagating this backwards:

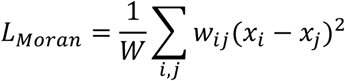

where, *L_moran_* is Moran’s loss, which quantifies spatial smoothness or autocorrelation; *W* is the total sum of all spatial weights; *w_ij_* is the spatial weight between locations *i* and *j*, based on proximity; and *x_i_* are feature values (e.g., gene expression, model outputs) at locations *i* and *j*.

To mitigate the issue of class imbalance in the training data, we applied weighted loss functions, ensuring balanced representation and learning across all classes:

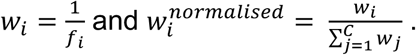

*f_i_* represents the frequency of class *i* in the training set. *w_i_* is the initial weight assigned to class *i* and 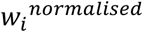 is the weight after normalisation. The normalisation step ensures that the sum of all weights 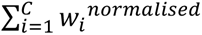 equals the number of classes *C*, maintaining balance in the loss function.

### GNN training and validation

Full-graph training was conducted to use the complete structural information present. When training, we used the entire graph by masking out the other non-tumour cells before loss back-propagation. Therefore, the masked out non-tumour cells were used for loss calculation but they did not influence the model as they were not used in the training. This improved model training compared to a more standard approach of using subgraphs.

GNN training can be approached using either transductive or inductive learning. In transductive learning, node classification is performed within a single graph, where some nodes are masked during training and used for prediction. Inductive learning, by contrast, involves training on subsets of a graph or entirely separate graphs, aiming to generalise the learned representations to previously unseen graphs. We reported the results for both approaches.

Spatial cross-validation was conducted using 10 spatial splits. When using the Xenium dataset, inductive training was conducted by dividing the slide into 10 spatial splits, while transductive training involved randomly masking nodes, keeping 10% of the nodes within a test set (5,776 tumour nodes). For the MERFISH dataset, which included 61 slides, inductive training was performed by splitting the data based on individual samples, whereas transductive training again involved random node masking across the entire dataset.

The models’ performances were evaluated using F1 scores and ROC-AUC for classification, and mean squared error for regression.

### GNN explanation

GNNExplainer was used for edge explanation. The main goal of GNNExplainer is to find a subset of the graph that maximises the prediction probability for a given target. This helps in understanding which parts of the input graph are most relevant to the decision made by the GNN. GNNExplainer learns a mask over the edge features, and uses gradient-based optimisation to update the masks. The loss function combines the prediction loss (which ensures that the masked subgraph results in the same prediction as the original) and a regularisation term (which controls the complexity and sparsity of the mask). We compared the explanations for each class to explanations generated from randomly shuffled nodes to obtain a p-value for each edge explanation.

Nodes were explained using integrated gradients, an approach which assigns an importance score to each node in a graph by measuring how adjusting that input node feature from a baseline to its actual value changes the model’s prediction:

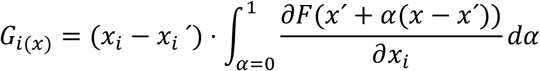

where, *i* is the feature; *x* is the input; *x*’ is the baseline; and *⍺* is the interpolation constant to perturb features by. The definite integral is not always numerically possible so a numerical approximation is calculated instead.

### Spatial regression modelling

Prior to spatial regression, we removed variables with high VIF and autocorrelation scores. We tested for additive effects using the spatial random forest spatialRF R package. Unless otherwise stated, we used models as implemented in the PySAL Python package^90^. We implemented spatial error modelling (SEM), an extension of linear regression that incorporates a spatially structured error term to model spatial autocorrelation in the residuals, thereby capturing spatial dependencies that would otherwise violate standard regression assumptions. The basic form of the spatial error model (SEM) can be expressed as:

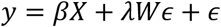

where *y* represents EMT labels for each observation; *X* contains the predictors for the regression model; *β* is the coefficient that represent the effect of each predictor; *W* is the spatial weights matrix (spatial relationships among observations); *λ* is the spatial autoregressive coefficient for the error term, which captures the strength of the spatial dependence in the error terms; and *ε* represents independent and identically distributed error terms, assuming a normal distribution *ε*∼*N*(0, *σ*^2^).

We applied SEM to various subsets of the Xenium data, including ductal carcinoma in situ (DCIS) and invasive regions, to assess how spatial interactions change within these regions.

Geographically weighted regression (GWR) and multi-scale geographically weighted regression (msGWR) were also used to further interrogate the spatial relationships. These fit local regression models for each point, and therefore answer different questions to a SEM model. SEM is important for understanding overall variable importance, and variance captured. However, it may not fit the most appropriate model for each spatial point, considering spatial heterogeneity. Comparing the model fit of SEM and GWR models allows for an assessment of how much heterogeneity is present, and to visualise how relationships change over space. GWR and msGWR were also compared for spatial fit. To evaluate model performance, R^2^ was used to measure the proportion of variance explained, providing an indication of how well each model predicts EMT. The Bayesian

Information Criterion (BIC) was also used to assess model complexity and fit, with lower values indicating a better balance between goodness of fit and model complexity.

GWR is a spatial analysis method that enables the modelling of spatially varying relationships between dependent and independent variables by fitting a local regression model at each point in space. The GWR equation can be represented as:

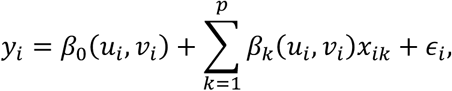

where *y_i_* is the EMT label at location *i*; *β*_0_(*u_i_*, *v_i_*) is the intercept term specific to location (*u_i_*, *v_i_*); *β_k_*(*u_i_*, *v_i_*) are the local regression coefficients for the *k*^th^ explanatory variable at location (*u_i_*, *v_i_*); *x_i_*_=_ are the explanatory variables at location *i*; *∈_i_* is the error term at location *i*; and *p* is the number of explanatory variables.

msGWR extends GWR by allowing each explanatory variable to have its own spatial bandwidth, which enables the modelling of multi-scale spatial relationships. The msGWR model can be represented as:

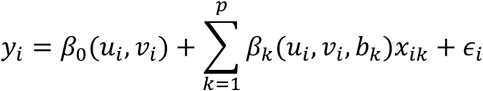

where *b_k_* is the unique spatial bandwidth for the *k*th variable, allowing the model to adapt to varying spatial scales.

GWR applies a single bandwidth across all variables, which may not capture multi-scale spatial processes accurately. In contrast, mxGWR, allows different bandwidths for each explanatory variable, provides a more flexible and specific understanding of spatial heterogeneity.

## Data availability

All data employed in this study is publicly available and was sourced from Zhang et al^36^ (MERFISH mouse motor cortex dataset) and Janesick et al^52^ (Xenium and Visium breast cancer dataset).

## Code availability

The spatial modelling framework developed in this study is available at: https://github.com/secrierlab/SPiCe.

## AUTHOR CONTRIBUTIONS

MS designed the study and supervised the analyses. EW performed all the analyses. CC critically assessed and refined the manuscript. All authors wrote and approved the manuscript.

## ACKNOWLEDGEMENTS

EW was supported by a studentship award from the Health Data Research UK-The Alan Turing Institute Wellcome PhD Programme in Health Data Science (218529/Z/19/Z). MS and CC were supported by a UKRI Future Leaders Fellowship (MR/T042184/1, MR/Y034031/1). Work in MS’s lab was supported by a BBSRC equipment grant (BB/R01356X/1) and a Wellcome Institutional Strategic Support Fund (204841/Z/16/Z).

## ETHICS DECLARATION

This study employs only publicly available data. All data comply with ethical regulations, with approval and informed consent for collection and sharing already obtained by the relevant consortia.

