## Supplementary Material for "Integrative Spatial Modelling of Cellular Plasticity using Graph Neural Networks and Geostatistics"

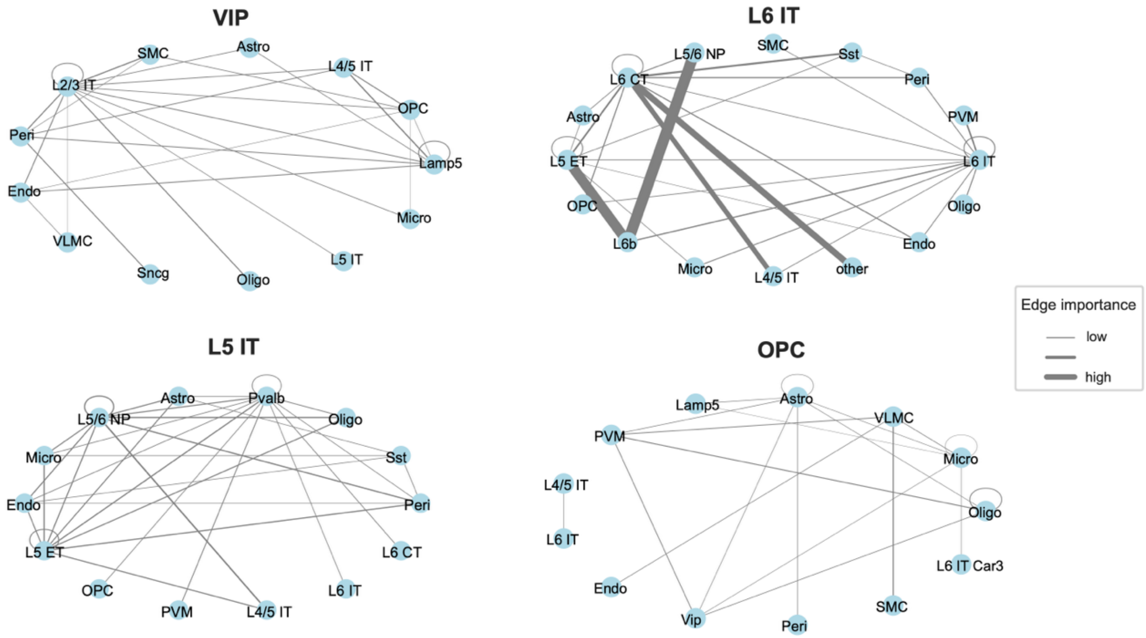


**Supplementary Figure 1. Edge importance graphs resulting from GNN modelling of the MERFISH mouse cortex dataset**. The cellular networks predictive of four selected types of cells (VIP, L6 IT, L5 IT and OPC) are shown.

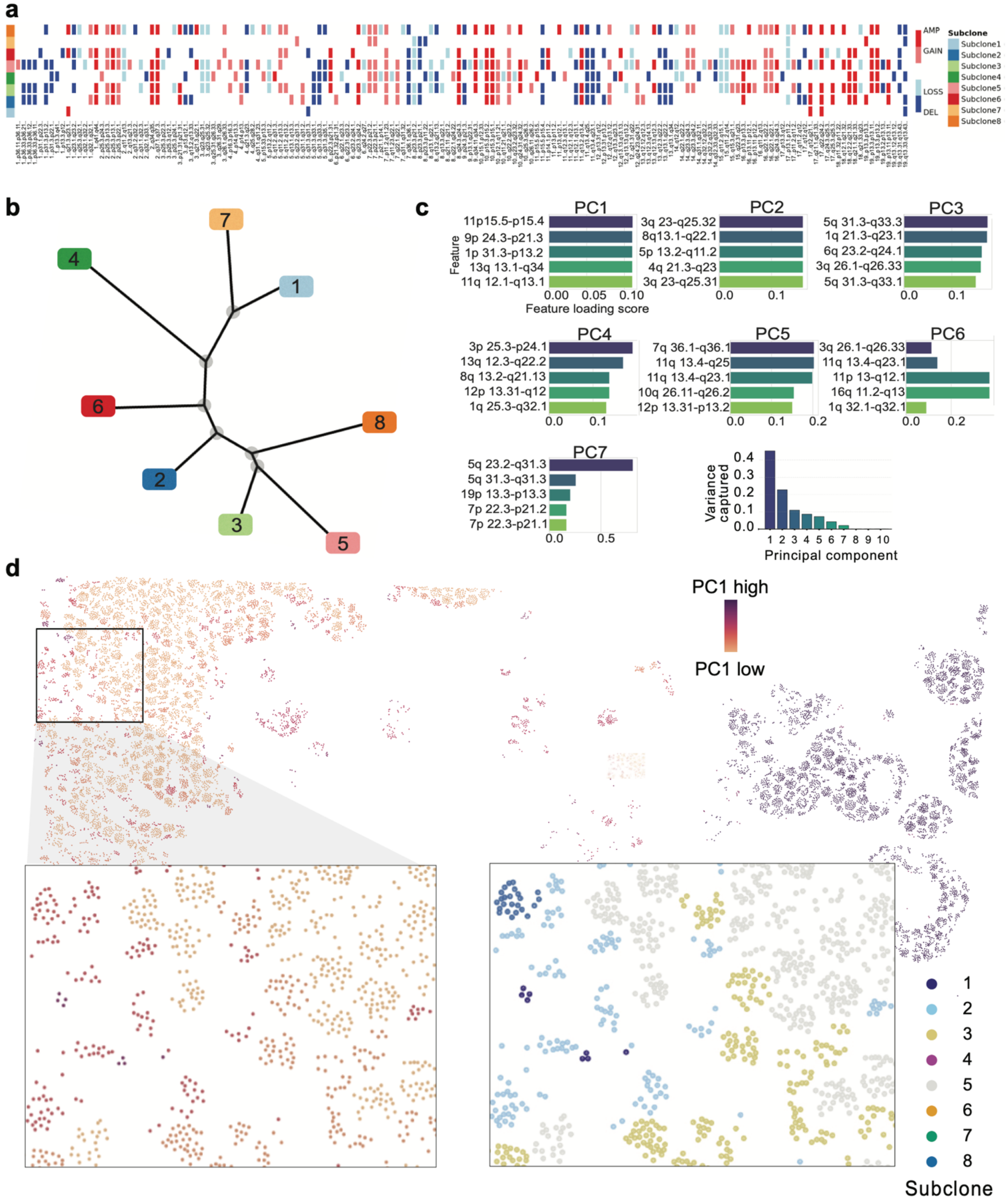


**Supplementary Figure 2. Tumour subclonality in the Xenium breast cancer dataset. a.** Copy number alterations across tumour subclones identified by SCEVAN in the Xenium breast cancer dataset. AMP = amplification; DEL = deletion. **b.** Clonal tree inferred by SCEVAN, highlighting subclones present in the tumour slide. **c.** Clonal alterations linked to the first seven principal components and the variance captured for each principal component. **d.** Principal component 1 visualised across the slide, with a zoomed in section of the slide with subclones identified.

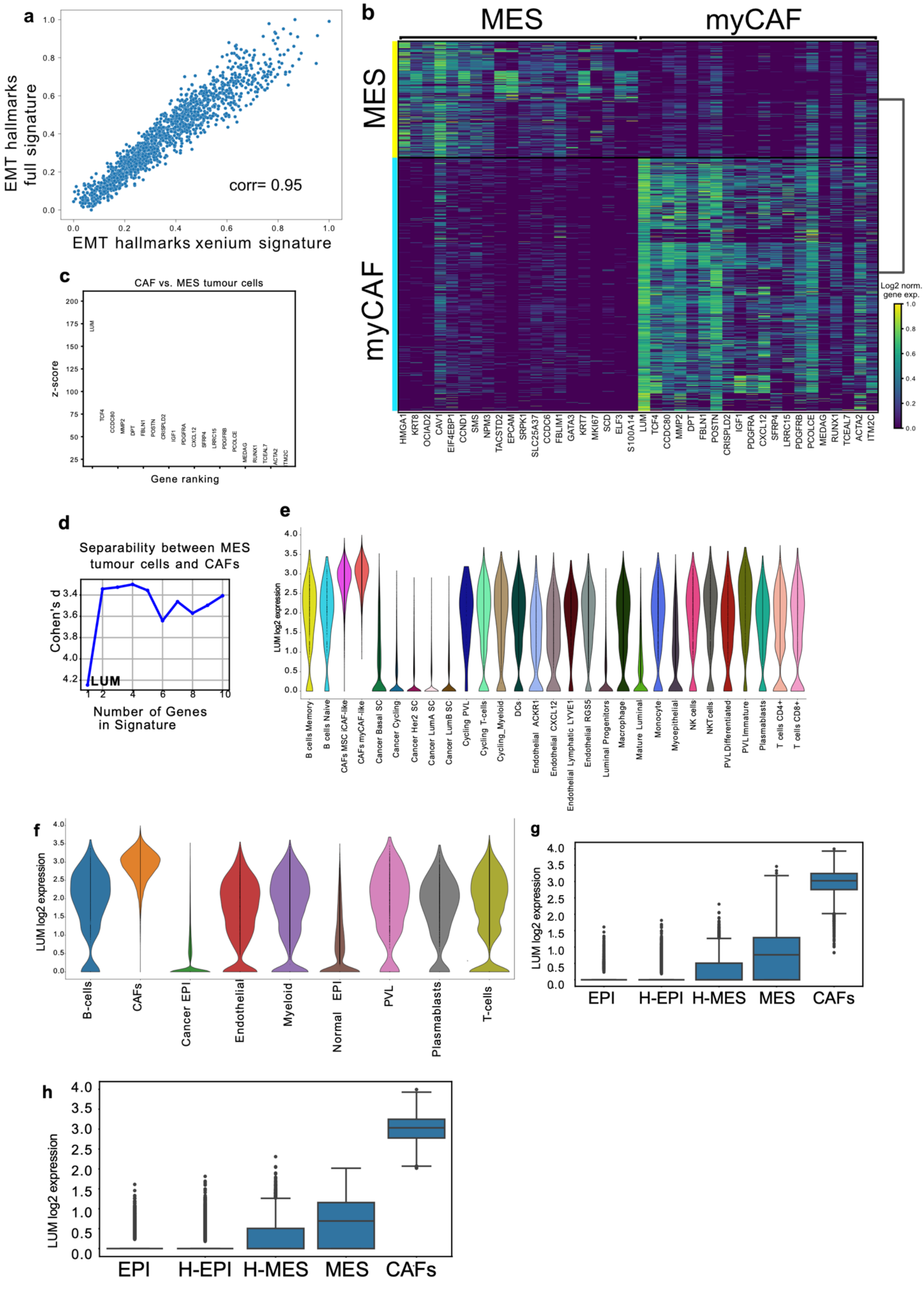


**Supplementary Figure 3. Validation of EMT signatures and discrimination between mesenchymal cancer cells and CAFs. a.** Correlation of EMT hallmark signature scores computed from the full gene panel (Visium) and the reduced Xenium panel (the Pearson correlation coefficient is displayed). **b.** Heat map of differentially expressed genes distinguishing myCAFs from mesenchymal-like tumour cells in the reference scRNA-seq breast cancer dataset. **c.** The top 10 discriminatory genes between myCAFs and mesenchymal tumour cells. *LUM* is significantly more discriminatory than other genes. **d.** Comparison of separability of cell types using Cohen's d score using *LUM* gene alone versus combinations of the next-best marker genes. *LUM* alone shows best separability (highest Cohen's d). **e.** Distribution of *LUM* expression across minor cell types in the scRNA-seq dataset. **f.** Distribution of *LUM* expression across major cell types in the scRNA-seq dataset. Expression is notably higher in myCAFs and iCAFs. **g.** *LUM* expression across cancer cell subsets and CAFs in the Xenium dataset prior to filtering. Expression is notably higher in CAFs, but some MES cancer cells show high expression too. **h.** *LUM* expression across cancer cell subsets and CAFs after filtering based on *LUM* expression. Good separability is achieved between CAFs and cancer cell subsets.

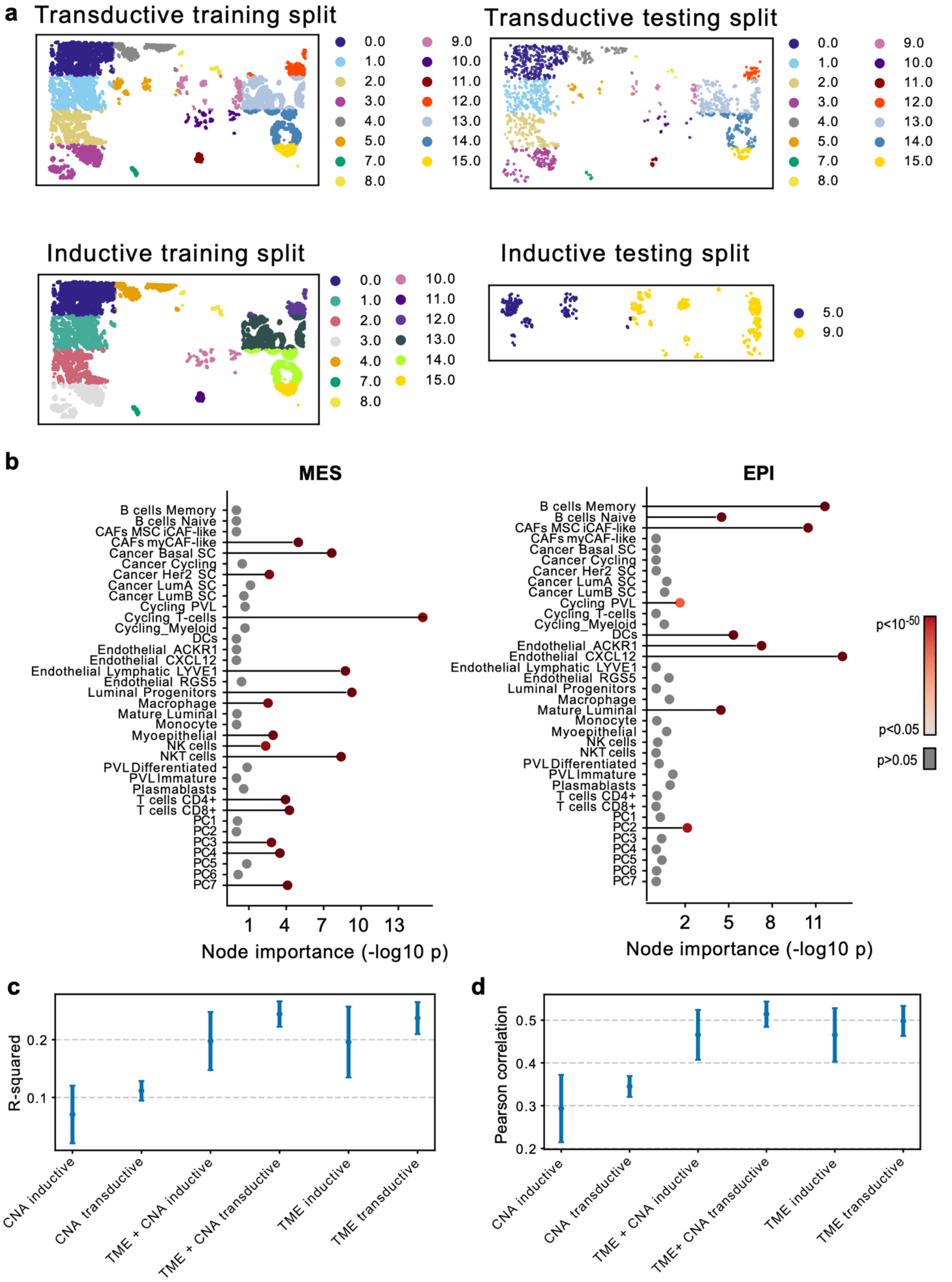


**Supplementary Figure 4. GNN modelling details and validation. a.** Illustration of train/test split using a transductive versus inductive split for GNN model training. In transductive training, train and test nodes come from the same graph, while in inductive training, the model is trained and evaluated on disjoint graphs. **b.** Simplified two-state EMT classification (epithelial versus mesenchymal) with node importance denoted by p-values. **c.** Predictive accuracy ($R^2$) of GNN regression models trained on continuous EMT scores using TME variables alone, CNAs alone, or the combination of CNA and TME variables, shown for both transductive and inductive learning settings. **d.** Predictive accuracy (Pearson coefficient) of GNN regression models trained on continuous EMT scores using TME variables alone, CNAs alone, or the combination of CNA and TME variables, shown for both transductive and inductive learning settings.

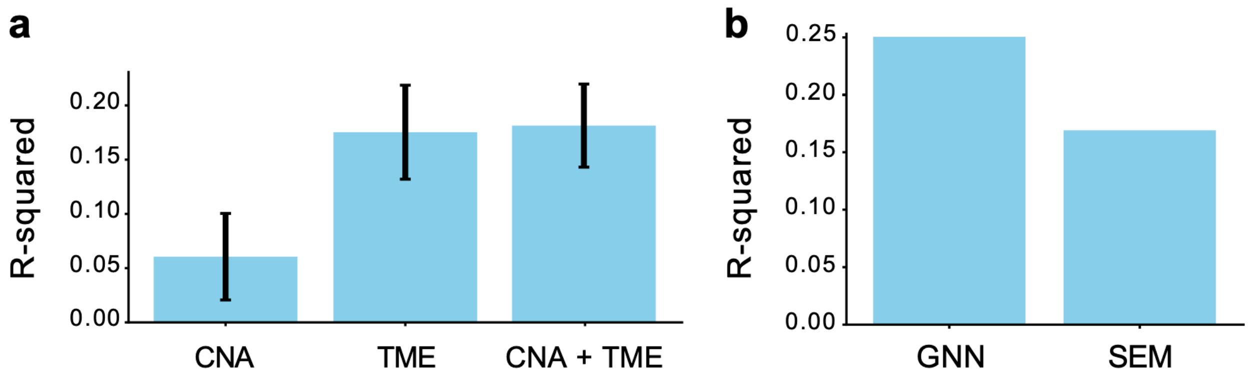


**Supplementary Figure 5. Performance comparison of spatial regression and GNN models. a.** Comparison of $R^2$ values from SEM models fit with TME and CNA predictors. **b.** Comparison of $R^2$ values between SEM and GNN regression approaches.
